# Unravelling the Maturation Pathway of a Eukaryotic Virus through Cryo-EM

**DOI:** 10.1101/2024.08.31.610394

**Authors:** Roger Castells-Graells, Emma L. Hesketh, Tsutomu Matsui, John E. Johnson, Neil A. Ranson, David M. Lawson, George P. Lomonossoff

**Affiliations:** Department of Biochemistry and Metabolism, John Innes Centre, Norwich Research Park, UK; UCLA-DOE Institute for Genomics and Proteomics, Los Angeles, CA, USA; Astbury Centre for Structural Molecular Biology, School of Molecular & Cellular Biology, Faculty of Biological Sciences, University of Leeds, Leeds, UK; Leicester Institute of Structural & Chemical Biology and Department of Molecular & Cell Biology, University of Leicester, Leicester, UK; Stanford Synchrotron Radiation Lightsource, SLAC National Accelerator Laboratory, Menlo Park, CA, USA; Department of Integrative Structural and Computational Biology, The Scripps Research Institute, La Jolla, CA, USA

**Keywords:** Virus maturation, eukaryotic virus, *Nudaurelia capensis* omega virus, virus-like particle

## Abstract

The importance of virus maturation has been appreciated for nearly 70 years ^1^ as it provides models for large-scale protein reorganization resulting in functional activation as well as being a target for antiviral therapies ^2^. However, a detailed description of the pathway from the initial assembly product (procapsid) to the mature, infectious particle (virion) has been elusive. This is due to the “in cell” nature of the natural process, the 2- state behavior of maturation (no detectable intermediates) in some viruses *in vitro* ^3^ and heterogeneous populations of particle intermediates that are only partially matured in other systems ^4^. The non-enveloped, T=4, ssRNA-containing, *Nudaurelia capensis* omega virus (NωV), is a highly accessible model system that exemplifies the maturation process of a eukaryotic virus. During maturation the particle shrinks in outer diameter from 482 Å (pH 7.5) to 428 Å (pH 5.0). It is possible to mimic the maturation process *in vitro* by lowering the pH of a population of procapsids produced in heterologous systems^5^. Indeed, by controlling the pH *in vitro* it is possible to produce homogenous populations of intermediate NωV virus-like particles (VLPs) that occur too fleetingly to be observed *in vivo* ^6^.

Here we report structural models, based on cryo-electron microscopy (cryo-EM), of five intermediates in the NωV maturation process. The structures of the intermediate particles reveal unique, quaternary position-dependent trajectories and refolding of subunit N and C-terminal regions, including the formation of the autocatalytic cleavage site at N570. The detailed structures reported here, coupled with previously determined structures of the procapsids and mature particles, allows the maturation pathway to be described in detail for the first time for a eukaryotic virus.

## INTRODUCTION

NωV is a T=4, ssRNA-containing alphatetravirus infecting pine emperor moth larvae in South Africa ^7^ and is a highly accessible model system with functionalities generally observed in eukaryotic virus maturation ^6^. Like some non-enveloped human viruses, its infection processes involve gene delivery through puncturing the plasma membrane with particle associated lytic peptides that are generated from the subunits during maturation ^8^. The icosahedral asymmetric unit (IASU) of NωV is formed by four copies of a 644-amino acid protein, each located in a distinct quaternary environment. Subunit A is adjacent to the icosahedral 5-fold axes and forms pentamer units with its symmetry equivalents; subunits B, C, and D are adjacent to the icosahedral 2-fold axes and form quasi-6-fold units with their symmetry equivalents (Fig. 1a). During maturation the particle shrinks in outer diameter from 482 Å (pH 7.5) to 428 Å (pH 5.0) as electrostatic repulsion is diminished by apoptotic-induced pH reduction in infected cells ^9^. Indeed, by controlling the pH *in vitro* ^5^ it is possible to sustain homogenous populations of intermediate NωV virus-like particles (VLPs), although such populations are too fleeting to observe *in vivo* ^9^.

**Figure 1.**
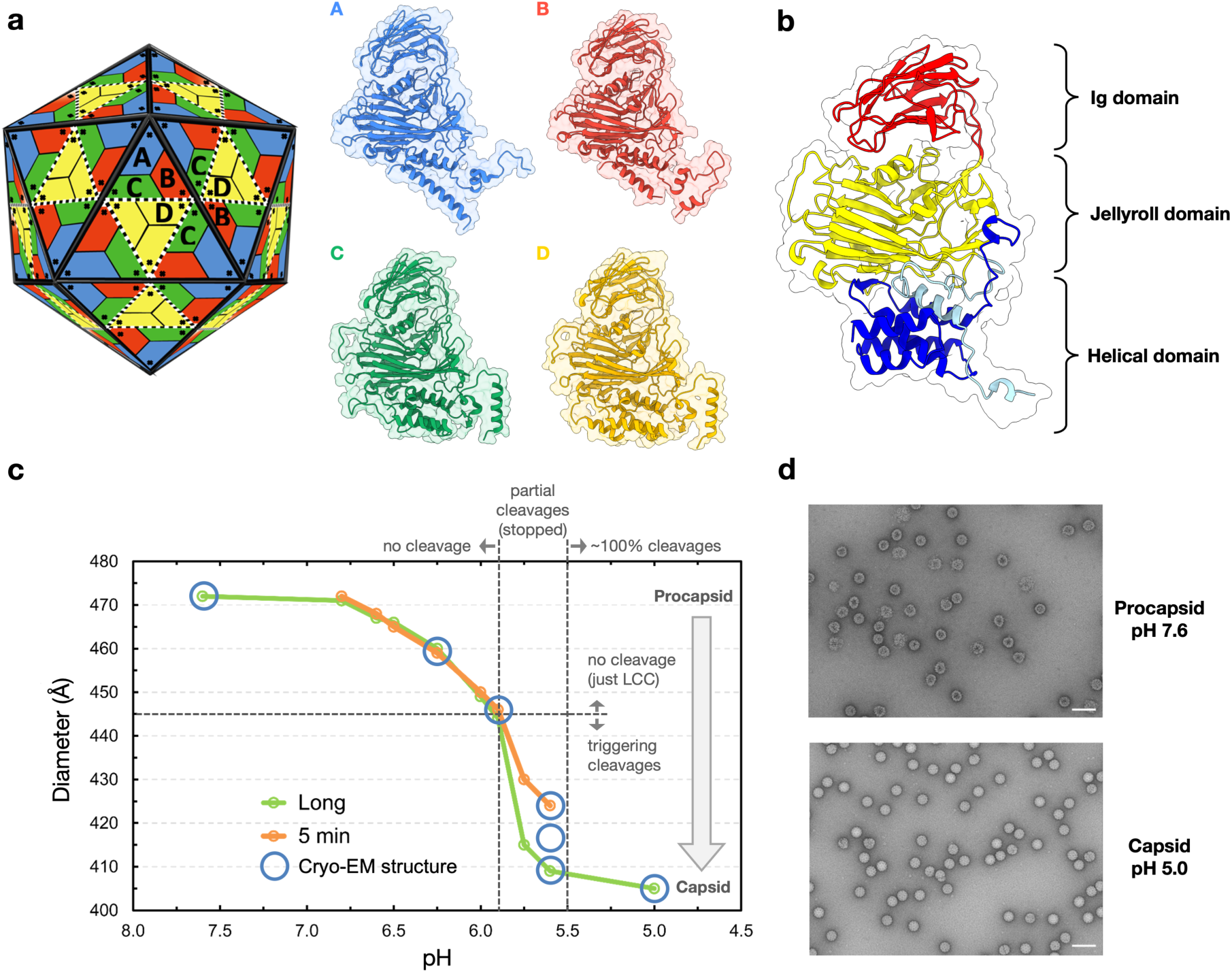
**(a)** Cartoon representation of the T=4 surface lattice of NωV showing the locations of the A, B, C and D subunits in the icosahedral asymmetric unit (IASU) of the particle. A subunits form pentamers; B, C and D subunits form quasi hexamers at the icosahedral 2-fold axes. The corresponding subunit tertiary structures of the mature particle are color-coded to the right and viewed in the same orientation with their outside at the top and their interior at the bottom. **(b)** The A subunit at pH 6.25 is shown in greater detail along with the virus jellyroll in yellow (residues 123-275 and 418-529), the inserted Ig-like domain in red (residues 276-417) and the internal N-terminal helices (73-124) in light blue and C-terminal helices (529-640) in dark blue. At pH 7.6 and 6.25 the interior helical domains are the same in A, B, C, and D. **(c)** Titration curves for NωV VLPs, employing the particle diameter (D_max_) as the “indicator”, from Small-Angle X-ray Scattering (SAXS) measurements at different times and pH values. The pKa for the particle at equilibrium is approximately 6. The particle radii were determined after incubation for 5 minutes at the given pH (orange) and after 6-7 hours (green). At pH values below 6 there is a clear annealing time for the particle to reach the equilibrium radius. Positions where cryo-EM structures were determined after 6-7 hours of incubation are shown with blue circles. At pH 5.6 the sample was incubated for 5 minutes and then frozen. Three main particle populations were found in class averages, as anticipated, for 5 minutes, which is the time required to see the first cleavages and the particles do not reach that state uniformly. **(d)** NωV VLPs stained with 2% (w/v) UA. The procapsids (pH 7.6), top, are porous and fragile and of an approximate diameter of ∼480 Å. With the pH drop and transition to the capsid stage (pH 5.0), the particles become less porous, more homogeneous and of an approximate diameter of ∼420 Å. Scale bar of 100 nm.

A refined atomic model of authentic, infectious NωV virions was determined by X-ray crystallography at 2.8 Å resolution ^10^, while the structure of mature NωV VLPs (non-infectious particles assembled in heterologous subunit expression systems and lacking genomic RNA) expressed in plants was determined by cryo-EM at 2.7 Å resolution ^11^. The two structures are identical within experimental error with less than 0.5 Å rmsd for 2256 aligned residues in the IASU ^11^ establishing the structural authenticity of VLPs and their usefulness for the maturation studies described here.

The structures revealed a 428 Å particle formed by 240 subunits, each with a canonical viral jelly roll ^12^ forming the contiguous shell and a 100 residue Ig domain ^13^, inserted between strands βD and βE of the jelly roll, that is displayed on the surface of the particle (Fig. 1b). Helical domains, formed by residues 44-116 in the N-terminal region and 560-644 in the C-terminal region, are on the inside of the particle, adjacent to the RNA. The C-terminal domain harbors an autocatalytic cleavage site (N570-F571) in the mature particle ^10^, cleavage of which liberates residues 571-644 in all four subunits. This C-terminal polypeptide remains closely associated with the particle, but cleavage allows these residues to play the role of a host cell membrane lytic peptide in the A subunits ^14^ and a particle stabilizing role in the C and D subunits ^10^. E103, on the same chain as the cleavage site, is a catalytic residue required for cleavage to occur ^15^. The formation of the active site and associated cleavage only occur at pH 5.6 and below. The inter-subunit surface is dominated by acidic residues ^16^ that are neutral below pH 5.0, the condition of crystallization. The structure of the procapsid, expressed in plants and purified at pH 7.6, has also been determined to 6.6 Å ^11^. From these prior studies, the start and finish points for the maturation process are largely known. What has been lacking is an understanding of the tertiary and quaternary structural rearrangements that occur in intermediates along the pathway to maturation.

## RESULTS

### Structures and dynamics of five NωV maturation intermediates

NωV VLPs have been the subject of numerous biochemical and biophysical studies ^6,17–21^ and display a reproducible and stable size reduction when the pH is adjusted at intervals between 7.6 and 5.0 in vitro ^22^. The titration curve, with the particle radius used as the “indicator”, is shown in Figure 1c as well as the pH values where cryo-EM structures were determined in the present study. Particle dimensions at pH 7.6, 6.25 and 5.9 were confirmed by SAXS (Table S1) with samples drawn from the same pool placed on the grid for the cryo-EM experiment. Likewise, the time dependent behavior of the sample at pH 5.6, over a period of 5 minutes, was monitored by SAXS with samples drawn from the same pool placed on the cryo-EM grid. Cryo-EM structures of the baculovirus expressed VLP procapsid (pH 7.6) and particles at pH values of 6.25, 5.9 and 5.6 (3 different structures at this pH), are reported here for the first time.

Procapsid, pH 6.25 and 5.9 particles were at equilibrium and displayed homogeneous populations on the EM grid. Particles at pH 5.6, where cleavage is first detected by SDS gel electrophoresis, were frozen on the EM grids 5 minutes after changing the pH of the buffer from 7.6 to 5.6. Under these conditions particles on the same grid were classified into three different populations, resulting in three reconstructions designated 5.6 Large (L), 5.6 Medium (M), and 5.6 Small (S) at the resolutions indicated in Table S2. The ordering of these structures toward maturation was based on particle diameter (L:448 Å, M:442 Å and S:430 Å) and the overall buried surface area (BSA) in the IASU (L:27,500 Å^2^, M:34000 Å^2^ and S:46000 Å^2^). The radial reduction correlates well with the increasing BSA of subunit interactions, supporting the proposed ordering of conformational forms.

The jelly roll and Ig domains in each subunit (residues 117-531) move as rigid bodies, each following distinct trajectories as the pH is lowered. During the pH titration, quaternary motions of the subunits are governed by the changing electrostatic fields generated at their surface. At neutral pH there is strong subunit repulsion by the large negative charge associated with the ionized inter subunit surface acidic residues, and this decreases as acidic groups become increasingly protonated at lower pH ^22^. Figure 2 shows the particles, IASUs and the positions tracked by the A subunit at pH 7.6, 6.25, 5.9 and multiple positions at pH 5.6. The motions of the A subunits follow that of a left-handed screw as they rotate counterclockwise about the 5-fold axes (viewed from the particle exterior) and move to lower radius. The trajectory of subunit B is similar to A and also twists significantly during its inward course, while C and D track a more linear radial path from 241 Å to 214 Å as the pH is lowered (Movies S1-6).

**Figure 2.**
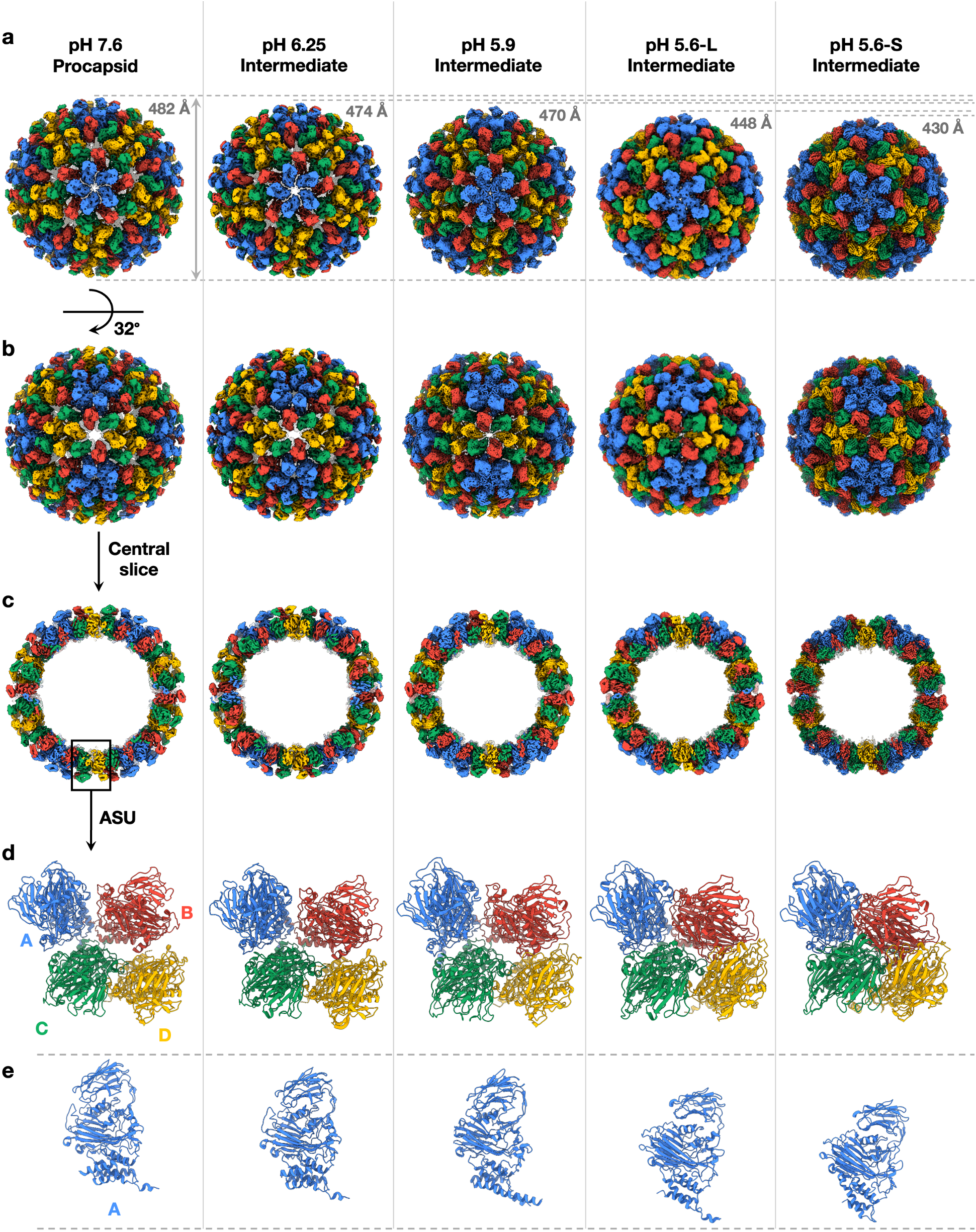
**(a)** NωV VLPs viewed down a 5-fold axis at pH values 7.6, 6.25, 5.9, 5.6 (large) and 5.6 (small). The particle diameters from left to right are 482 Å, 474 Å, 470 Å, 448 Å, 430Å. Subunits are color coded A (blue), B (red), C (green) and D (yellow). **(b)** NωV VLPs viewed down a quasi-6-fold (icosahedral 2-fold) axis. **(c)** View of the central slice of NωV VLPs. **(d)** The IASU viewed from outside the particle with the same subunit color coding and showing the change in subunit contacts at the same pH values as in a. BSA for the IASU at these pH values are 14500 Å^2^, 16500 Å^2^, 21000 Å^2^, 27,000 Å^2^ and 46,000 Å^2^. **(c)** The A subunit viewed from the side at the same pH values as a, b, c and d showing changes in radius, subunit orientation and internal helical domain.

During maturation, local tertiary structural changes in the four subunits are confined to residues 42-116 at the N-terminal region and 560 to 644 at the C terminal region. Residues prior to 42 are never visible in the density of any of the structures, and these 41 residues contain 13 R and K basic amino acids. It is assumed that residues in this region associate with the RNA and allow the initial formation of the procapsid at neutral pH. As described previously ^11^ and below, only weak inter-subunit interactions are present at the procapsid stage. Thus, the procapsid subunits can be visualized as a collection of “balloons” (residues 42-644) tethered by strings (residues 1-41) to the core RNA, with “balloon” quaternary positions influenced by the electrostatic fields associated with the negatively charged surfaces of the A, B, C, and D subunits at neutral pH.

### First step in the maturation: from pH 7.6 to pH 6.25

The first step in the observed maturation occurs between pH values 7.6 (procapsid) and 6.25 (the first intermediate structure). These two points in maturation provide insights into the likely state of the assembling protein and its unassembled tertiary structure (Figure 2). Unlike later in maturation, the internal helical C-terminal structures (residues 548-642) of A, B, C, and D are closely similar to each other as the diameter decreases from 482 Å to 474 Å (Figure 3). This observation is consistent with the notion that initial assembly occurs with all subunits virtually identical in structure, probably resembling their solution structure in the unassembled state. These residues form five-helix bundles that translate and rotate as a rigid unit with the jellyroll/Ig domain between these pH values. They form the principal points of subunit interaction that stabilize the early-stage particle. At pH 7.6 and 6.25 the particle is stabilized by subunit dimers and trimers. The A2-B1 and C1-D1 dimer BSA values are equivalent and have the largest value of ∼2800 Å^2^, with 35 of the 42 contact residues being in common for these unique interfaces, consistent with dimers representing the solution state and initial assembly state of the subunits. The A, B, C quasi-trimer is virtually identical to the D, D, D icosahedral trimer (Figure 4). In addition to interactions between the helical domains the trimers are stabilized by a small domain swap between subunits related by 3-fold or quasi-3-fold symmetry at residues 74-80 of the N terminal visible region (residues 74-117) which is also part of this internal module leading to the next highest BSA at ∼1700 Å^2^ and 18 contact residues shared of a total of 25 at each of the four unique contacts. Procapsid and pH 6.25 particles provide novel insights into particle assembly as the tertiary structures for all of the subunits are closely similar and the quaternary structures have not yet been affected by maturation. The subunit tertiary structures of the procapsid and pH 6.25 particle demonstrate that the cleavage site (composed primarily by essential residues E103 and N570 on the same chain) is not formed. The separation of these residues is ∼16 Å in all four subunits (Figure 5). This separation is too large to enable the active site residues to catalyze cleavage prior to maturation, indicating that, as described below, proteolytic active site formation is dependent on the changes in quaternary structure at lower pH values.

**Figure 3.**
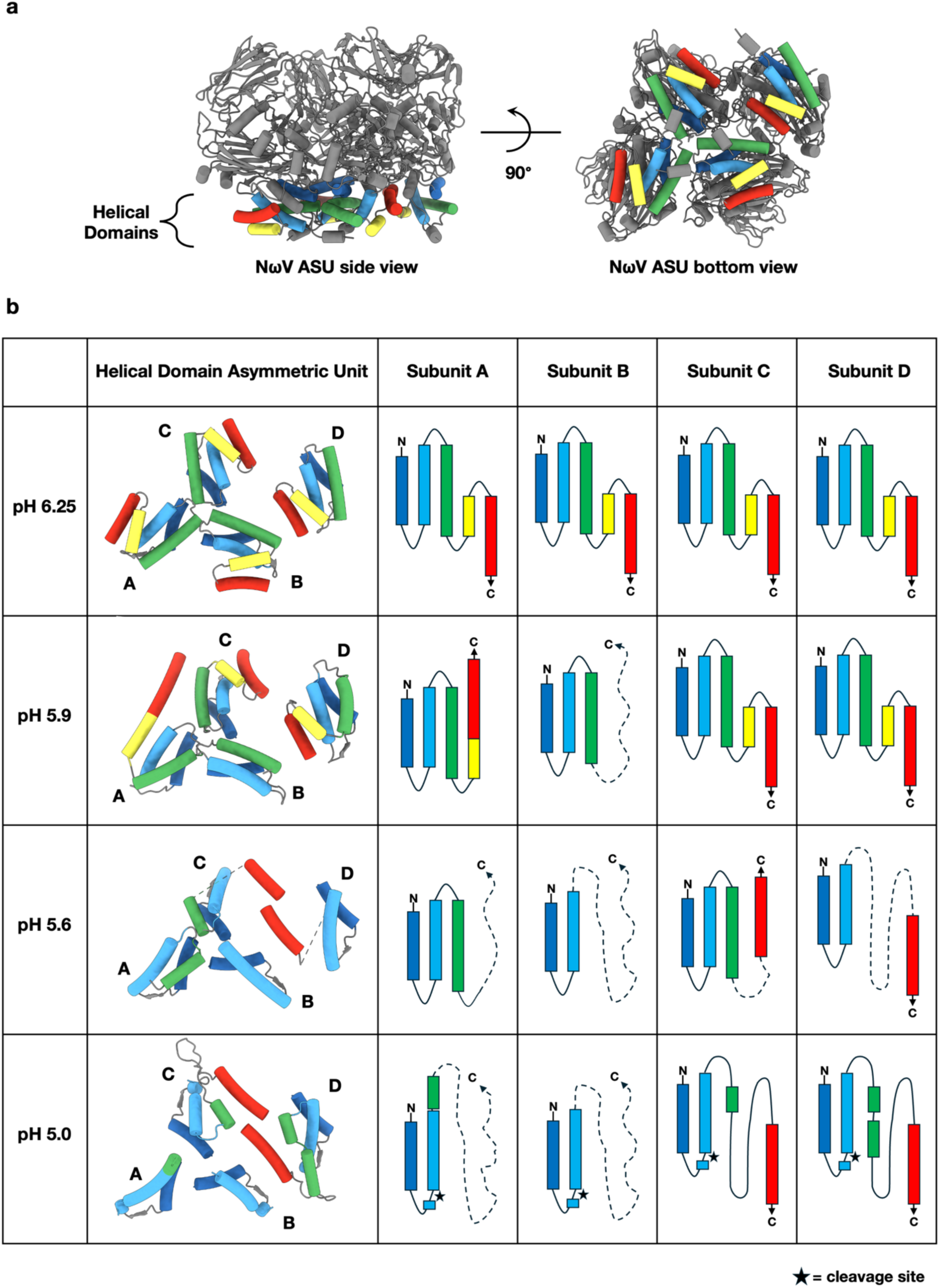
Schematic representation of the refolding of the internal helical domains shown for the four subunits in the IASU at 4 different pHs. **(a)** Side and bottom view of the IASU at pH 6.25 with the internal helical domains colored. **(b)** At pH 6.25 the internal helical domains of A, B, C and D have the same structure, with residues indicated for each helix. The five C terminal helices are color coded blue (548-560), turquoise (571-585), green (592-608), yellow (613-620) and red (627-643) in this panel. The color coding for the helices is maintained at the different pH values to show how they reorganize during maturation. At pH 5.9 the A subunit refolds with helices 4 and 5 fusing to form one long helix. The B subunit helical region begins to disorder with the last visible residue 608. C and D have essentially the same structures as they had at pH 6.25. At pH 5.6 more refolding has occurred and all four subunits are different. The A subunit now has the residues ordered that are nearly the same as those observed in the in the capsid (pH 5.0); 548-608. In B, the last observable residue is 590. Residues in C are continuously visible from 548-602 and 627-643 while in D the last observable residue is 642, but residues 589-626 are disordered. At pH 5.0 the particle is mature. The A subunits (cleaved residues 571-597) form the helical, membrane lytic module, the B subunits are disordered after 590 and C and D subunits are fully ordered to the end of the polypeptide and form the molecule switches that stabilize the particle.

**Figure 4.**
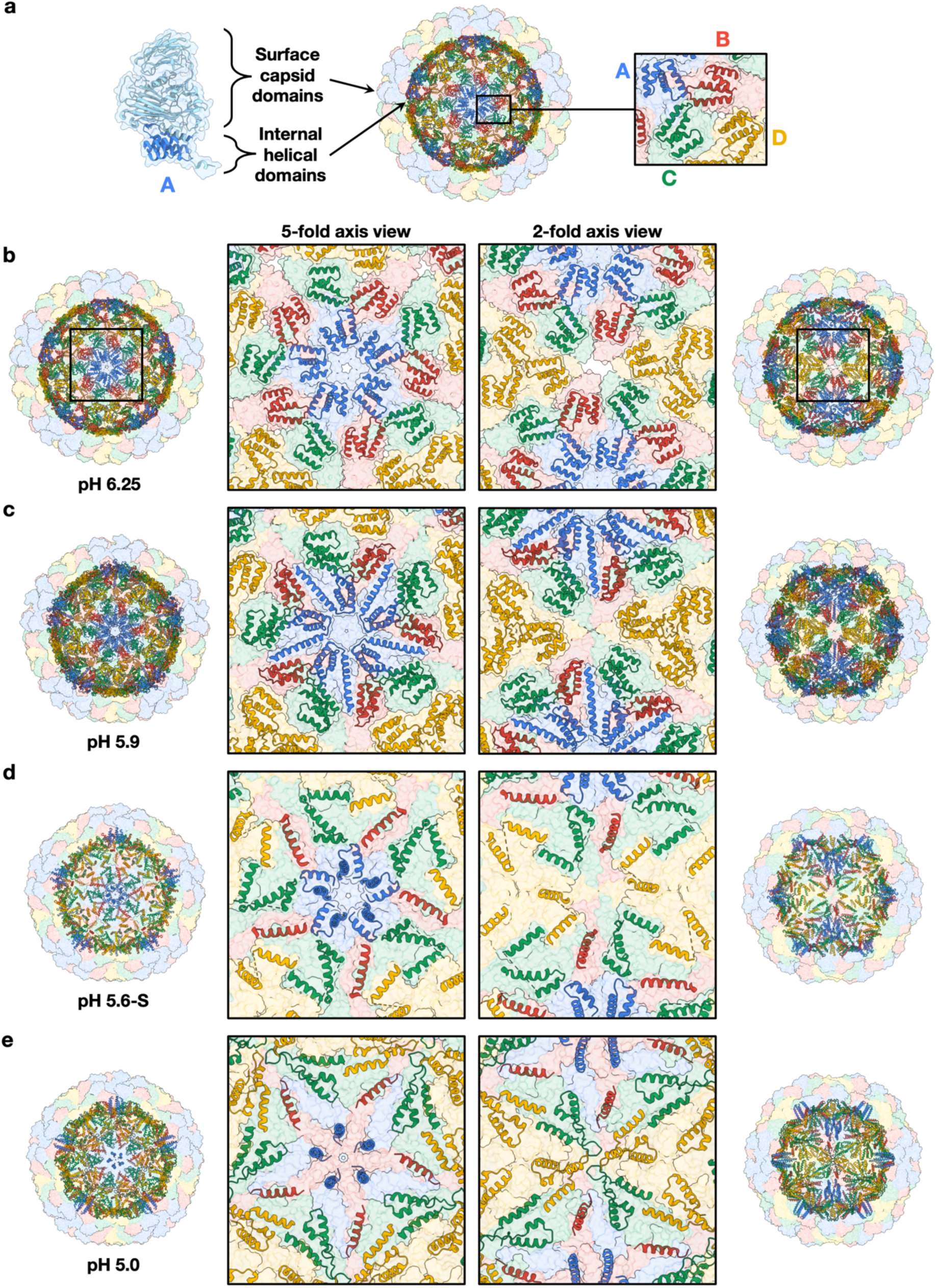
The quaternary structure environments of the icosahedral asymmetric units for the different intermediates shown in Figure 3. **(a)** A subunit at pH 6.25 next to a transparent surface view of a NωV particle with colored cartoon representations of the internal helical domains. Color coding corresponds to A (blue), B (red), C (green), D (yellow) subunits. **(b)** Intermediate at pH 6.25, the C-terminal helical domains in the four subunit types are virtually identical. Particles and internal helical region (residues 571-644) viewed down the 5-fold and quasi-6-fold (icosahedral 2-fold) particle axis. **(c)** Intermediate at pH 5.9 showing the extended C terminal helix in the A subunits invading the space occupied by the B and C subunits. **(d)** Intermediate at pH 5.6. Helices in the A subunits are poised to refold again to form the helical bundle, membrane-lytic, module. The C and D subunit terminal helices (626-641) are positioned to form the molecular switch at the quasi-2-fold axes. **(e)** Final disposition of helices in the mature capsid (pH 5.0). The autocatalytic cleavage has occurred, covalently releasing A helices in the bundle that are ordered from 571 to 600. The remaining residues in the A helices are disordered and may be attached to the RNA. C and D terminal helices are also covalently independent and form the molecular switches adjacent to the quasi-2-fold axes. The helical structures in C and D are closely similar to each other, while the A and B subunits show a different structure for the same residues.

**Figure 5.**
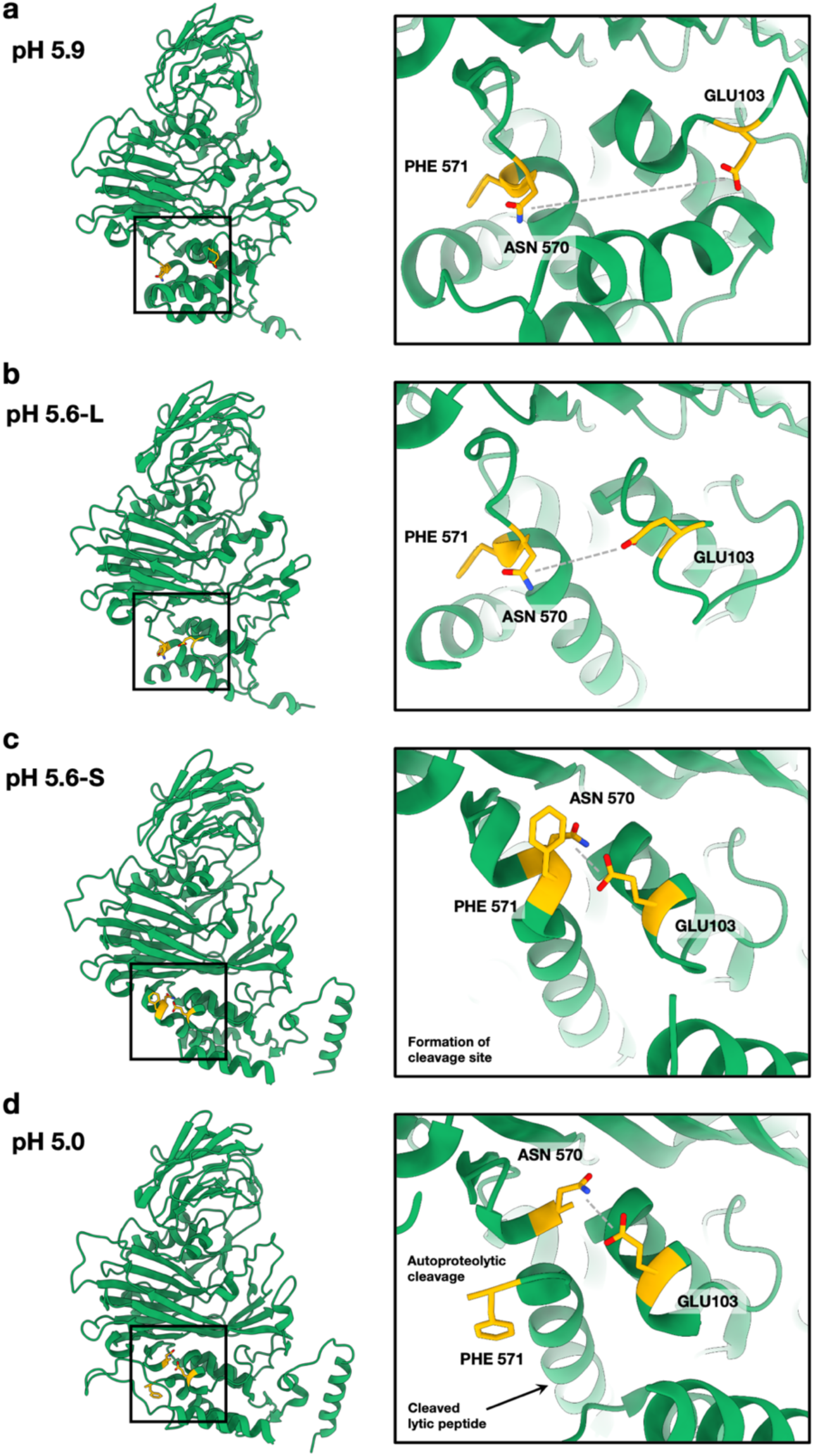
The motion of Glu 103 (the required catalytic residue) and Asn 570 (the residue where cleavage occurs) as a function of the maturation intermediates. **(a-b)** Before the formation of the cleavage site the distance between Glu 103 and Asn 570 is ∼9-16 Å. **(c)** The cleavage site is formed, the distance between Glu 103 and Asn 570 is ∼4 Å and the autoproteolysis is initiated. **(d)** The autoproteolytic cleavage is completed and residues 571-644 are liberated in all four subunits.

### Refolding of the internal helical domains and formation of the cleavage site below pH 6.25

For the virus to attain its final infectious form, it requires that the tertiary structures of the C-termini change significantly, and differently in different subunits, as the pH is lowered below 6.25 (Figure 3). In the procapsid, this region is essentially the same in all four subunits and folds into five helices, which we shall refer to as helices 1-5. Between pH 6.25 and 5.9 there is a dramatic change in the C-terminal residues of the A subunit where residues 613 to 641, originally forming the last two helices (613-620 and 627-641) of the 5-helix bundle, refold into a single long helix that is now part of a 4-helix bundle clustered about the 5-fold axes but with the long helix also invading space occupied by an adjacent B subunit (Figure 4). This is reflected by a 30% increase in the BSA for A2-B1 compared to C1-D1, diverging from values that were equivalent at pH 6.25. These changes are the first events in the assembly of the membrane lytic module in the A subunits at the pentamers. The B subunit helical domain is significantly disordered at pH 5.9 with helices 1, 2 and 3 retaining visible structure and residues 608-642 invisible. The internal helical domains of the C and D subunits do not significantly change structure between pH 6.25 and 5.9. The buried surface areas within the trimers remains low at ∼2000 Å^2^ indicating little increase in their contribution to particle stability. There is a small change in formation of the cleavage site with the average separation of E103 and N570 at pH 5.9, reducing to 14 Å (Figure 5).

A critical particle transition for infection occurs at pH 5.6 where the first evidence of limited cleavage is observed. The three structures determined at this pH (L, M and S) provide significant insight regarding positioning of the C-terminal helices in A, C and D subunits and their distinct roles for lysis of the target cell plasma membrane (A subunits) and particle stability (C and D subunits), as well as formation of the cleavage active site (E103-N570). At pH 5.6 the long helix in A becomes disordered, leaving residues 548-614 ordered (forming helices 1-3) and the remainder disordered (i.e. invisible in density maps). Subunit B has only helices 1 and 2 ordered, subunit C has helices 1-3 (548-602) and 5 (627-642) ordered and subunit D has helices 1 and 2 ordered and a disconnected well-ordered helix 5 (residues 627-642). An appropriate metric for relating the 3 transition intermediate structures determined at pH 5.6 is the comparison of the A2-B1 contact with the C1-D1 contact. These contacts occur at quasi-equivalent 2-folds that are closely similar at pH 7.6, 6.25, and 5.9 but which diverge at pH 5.6 into what are called the “bent” A2-B1 contact and the “flat” C1-D1 contact in the fully mature particle. Comparing equivalent residue contacts at these two quasi-2-fold axes demonstrates that intermediate 5.6-L has 23 of a total of 40 contacts equivalent (compared to 35 of 42 at pH 6.25), 5.6-M has 20 of 40, and 5.6-S has 19 of 50. In addition, while L and M have similar buried surface areas in the two contacts, S shows that C1-D1 has almost three times the buried surface compared to A2-B1. This is owing to the insertion of helix 5 (626-642) in subunit D creating a larger BSA at the flat contact. Following the motions of all the C-terminal helices (548-642) in D demonstrates that, prior to cleavage, disorder between 589 and 626 is required for insertion of the 626-642 ordered helix. As described below, following cleavage between 570 and 571, all the residues from 585 to 642 are ordered in both C and D subunits, demonstrating the critical role of cleavage in making the independent C-terminal regions of these subunits sufficiently flexible to insert the helices (molecular switches) and create the final stable particle. A second metric of the maturation process is the BSA at the quasi (ABC) and icosahedral (DDD) 3-fold axes. These move from 3000 (pH 6.25) to 4000 (pH 5.9) to nearly 6000 Å^2^ in pH 5.6 L, M, and S, and are similar to each other in the progression as the particle condenses. While minimally active, cleavage site formation does occur at this pH with the average distance between E103 and N570 in the 4 subunits being ∼10 Å for L and M, followed by a dramatic reduction to ∼4 Å in S. The acrobatic motions of E103 and N570 between 5.6 L and S that bring the residues into juxtaposition are shown in the movie of the active site formation (Figure 5 and Movie S7).

### Autoproteolysis and formation of the lytic peptide in the last maturation step

At pH 5 the VLP capsid is fully formed, and the subunits are cleaved at residue 570 generating a particle whose structure is identical to the infectious virion. Following the cleavage site, the A subunit forms a single long helix (574-596) composed of original helices 2 and 3, and these are near the 5-fold axes. These covalently independent helices have been shown to be a membrane lytic module that escapes the particle when it is in proximity to a target membrane. The long and disordered region (597-644), with a very basic C-terminal region (10 of the 25 terminal residues are R and K), makes it plausible that this module may interact with and deliver genomic RNA into the cell in concert with lysis. C-terminal regions (571-641) of the C and D subunits are fully ordered and modeled in the mature particle. Helices 627-642 are molecular switches that buttress the C and D subunits forming the flat contact. They conform closely to the expected quasi-2-fold symmetry. These molecular switches are disordered in the A2-B1 quasi-2-fold contact allowing the subunits to pivot about a hinge (bent contact) creating a totally different set of subunit interactions below the hinge when compared with C and D.

## DISCUSSION

The structures reported here demonstrate a remarkable modularity in the subunit sequences. Residues 117-531 (jellyroll with inserted Ig domain) behave throughout as a rigid structure that is superimposable for the A, B, C, and D subunits at all pH values investigated. The novelty of the quasi-equivalent positions is governed by electrostatic forces between these rigid structures and by the N and C terminal regions (45-117 and 535-644) of the subunits. These residues form the different functionalities (lytic polypeptide A subunits; molecular switches C and D subunits) described and yet all four subunits form identical autocatalytic cleavage sites with the exceptional precision required for the chemistry. This is achieved through a motion of greater than 10 Å (between procapsid and intermediate S at pH 5.6) that brings E103 into perfect position to function as a base that catalyzes the nucleophilic attack of the side chain of N570 on its main chain carbonyl carbon to form a cyclic imide that is then hydrolyzed and cleaved (Figure 5). The presented structures provide a wealth of detail about the dynamics of this programmed nanomachine, and may be used to train new models to aid in the design of novel protein assemblies, such as dynamic protein cages, using machine learning algorithms ^23^.

The choreography between particle stability and infectivity is controlled by large scale quaternary structural changes that dictate critical tertiary structure alterations at the N and C-terminal regions of the subunits. The precision in triggering these changes at the latest stage of infection with apoptotic-associated pH reduction and only then making released, infectious virions, is a wonder of viral evolution. While the present work has documented the relationship between these changes and has mapped them in detail, the cause-and-effect correlation remains elusive and the subject of further study, including by computational methods. The structural interplay described here is a microcosm of larger scale events in biology that commonly involve changing protein-protein interactions and chemical modifications to the associated players.

## MATERIALS AND METHODS

### Virus-like particles preparation and purification

*Spodoptera frugiperda* cells (Sf21 cell line) were grown in an incubator with shaking at 26°C and infected with recombinant baculoviruses when they reached a concentration of 2 to 3 x 10^6^ cells x ml^-1^. The baculovirus stocks were generated as per manufacturer’s instructions. Insect cells transiently expressing the coat protein of NωV were harvested at 3 to 5 days post infection. The infected insect cell culture was spun at 500 x g for 10 minutes at 11°C. The pelleted cells were resuspended in 25 ml of 50 mM Tris, 250 mM NaCl, pH 7.6 buffer for every 50 ml of initial cell culture. Nonidet P-40 (NP-40) was added to a final concentration of 0.5 % (v/v). Then the mixture was incubated for 15 minutes on ice and then spun for 10 minutes at 10,000 x g. The supernatant was transferred into the ultracentrifuge tube (UltraClear 25 x 89 mm) and then 3 ml of 30% (w/v) sucrose solution were layered at the bottom of the tube. The sucrose cushion was centrifuged at 166,880 x g (30,000 rpm) for 3 hours at 11°C. The supernatant was discarded and 400 μl of the extraction buffer were added to the pellet to resuspend it overnight at 4-8°C on a shaking platform. The resuspended pellet was spun at 12,000 x g for 30 minutes at 11°C. The clarified supernatant was transferred on top of a 10-40% (w/v) continuous sucrose gradient and it was centrifuged at 273,800 x g (40,000 rpm) for 1 hour and 15 minutes at 11°C. Gradient fractions containing the virus-like particles were identified by SDS-PAGE and concentrated using centrifugal filters (Amicon^®^, Merck) with a molecular weight cut-off (MWCO) of 100 kDa. The concentrated VLPs were stored at 4°C.

### SAXS analysis of VLPs samples

Small-Angle X-ray Scattering (SAXS) experiments were conducted at Beamline 4-2 (BL4-2) of the Stanford Synchrotron Radiation Lightsource (SSRL). The experimental setup is summarized in Table S1. Briefly, data were collected on a Pilatus3 X 1M detector (DECTRIS, Switzerland) with a 1.7-m sample-to-detector distance, and the beam energy of 11 keV (wavelength, λ = 1.127 Å) was used. The momentum transfer (scattering vector) q was defined as q = 4πsin(θ)/λ, where 2θ was the scattering angle. The q scale was calibrated by silver behenate powder diffraction ^24^. A series of VLPs samples (pH-titration series) and its equivalent buffer were placed on a 96-well plate of BL4-2 SAXS Autosampler, which was operated by the BL4-2 data acquisition program, *Blu-ICE* ^25^. A 1.5-mm-quartz capillary cell (Hampton Research, Aliso Viejo, CA) was maintained at 20 °C. The sample solution loaded to the cell was oscillated during exposures to alleviate radiation damages.

The 10 scattering images with 3 sec exposure were obtained from a 30 μl sample or buffer aliquots. The SAXSPipe, an automated SAXS data processing and analysis pipeline of the BL4-2 (https://www-ssrl.slac.stanford.edu/smb-saxs/content/documentation/saxspipe), was employed for automatic data processing and initial analysis. The background-subtracted data were then used for further analysis.

The size of the VLPs was evaluated by P(r) analysis using the program Gnom ^26^. To eliminate the effect of inter-particle interactions induced by the reduction of negative electrostatic charges at lowering the pH value ^16^, the q-range from 0.012 to 0.08 Å^-1^ was used for the analysis.

### Negative staining of VLPs samples

Grids for negative staining were prepared by applying 3 μl of sample (∼0.1 to 1 mg/ml) on to carbon-coated 400 mesh cooper grids (EM Resolutions). The grids were glow-discharged for 20 seconds at 10 mA (Leica EM ACE200) prior to applying the sample, Excess liquid was blotted away with filter paper and then the grid was stained with 2% (w/v) uranyl acetate (UA) for 30 seconds.

### Cryo-electron microscopy sample preparation and data collection

Cryo-EM grids were prepared by applying 3 μl of sample (∼0.2 to 0.4 mg/ml) to 400 mesh copper grids with a supporting carbon lacey film (Agar Scientific, UK) held on an automatic plunge freezer (Vitrobot Mk IV). The lacey carbon was coated with an ultra-thin carbon support film, less than 3 nm thick (Agar Scientific, UK). Prior to applying the sample, grids were glow-discharged for 30 seconds (easiGlow, Ted Pella). The samples were vitrified by flash-freezing in liquid ethane, cooled by liquid nitrogen.

Data were collected on an FEI Titan Krios EM at 300 kV (Astbury Biostructure Laboratory, University of Leeds). The exposures were recorded using the EPU automated acquisition software on a FEI Falcon III direct electron detector. Micrographs were collected with a final object sampling of 1.065 Å/pixel.

### Cryo-electron microscopy data processing and model building

Movie stacks were motion-corrected and dose-weighted using MOTIONCOR2 ^27^ (Fig. S1). CTF estimation was performed using GCTF ^28^ and particles were picked using RELION ^29,30^. Automatic particle picking was performed using 2D templates generated after an initial run without reference templates (Laplacian). Subsequent data processing was carried out using the RELION 2.1/3.0 pipeline ^29–31^ with the imposition of icosahedral symmetry for the 3D reconstructions (with the exception of the pH 5.6 intermediate datasets where symmetry expansion was employed, as described below). The pH 5.9 intermediate model at 3.92 Å resolution was generated first, starting from the previously published cryo-EM structure of the plant-expressed capsid VLPs (PDB entry 7ANM). The icosahedral asymmetric unit (IASU), comprised of four protein chains, was rigid body fitted to the sharpened map in Chimera ^32^. To expedite computation, for the subsequent steps, the IASU was visualised, manipulated and refined in the context of its eight nearest symmetry copies, denoted IASU8. The central IASU was edited using COOT ^33^ with reference to unsharpened and sharpened maps and refined using phenix.real_space_refine ^34^ against the latter. An updated IASU8 was generated from the central IASU after each refinement job. Validation of the final model was performed on the IASU using Molprobity ^35^ through the Phenix interface ^36^. The final pH 5.9 model (PDB entry 8A3C) was used as the starting point for generating the models of the pH 6.25 intermediate and procapsid, at resolutions of 4.80 and 4.88 Å, respectively, using similar protocols.

For the pH 5.6 intermediate, a single dataset was heterogeneous, so it was combined with a second dataset to ensure sufficient particles after multiple rounds of classification for meaningful reconstructions. Initially, 3D classes could be distinguished on the basis of size, although there was most likely a continuum of different sizes, with the smaller, more capsid-like classes refining to higher resolutions than the larger more “pH 5.9-like” classes. Focussing on the smallest class comprised of 15,977 particles, it was possible to achieve a resolution of 3.39 Å after postprocessing. A preliminary model was built as described for the other datasets starting from the plant-expressed capsid VLP structure (PDB entry 7ANM). However, the map was difficult to interpret in many places, despite the resolution. Given the heterogeneous nature of the data, we hypothesised that the particles might not be perfectly symmetrical and that by imposing icosahedral symmetry we were averaging the densities for particles in multiple conformational states leading to blurred maps. Thus, we resorted to using symmetry expansion within RELION to enable the heterogeneity to be explored. Briefly, the particles from the best 3D refinement job of the smallest class were expanded to C1 symmetry and further 3D classified without alignment into 10 classes using a soft mask covering just the IASU. For each of the three most populated classes (accounting for 36, 27 and 10% of the stack), particles were randomly split into half sets for the subsequent reconstruction and postprocessing steps, giving reconstructions to 3.39, 3.63 and 3.91 Å resolution, respectively. From these maps it was possible to build and refine three discrete versions of the IASU using the same protocols described above. Since the final reconstruction was performed using C1 symmetry, the assumption is that the particle does not obey strict icosahedral symmetry. However, to provide the approximate context for these IASU models, we used icosahedral symmetry operators to place them back into a full VLP protein shell. From these models, it was clear that the three reconstructions gave rise to particles with different diameters, which we describe as “small” (diameter ∼415 Å, resolution 3.39 Å), “medium” (diameter ∼425 Å, resolution, 3.63 Å) and “large” (diameter ∼430 Å, resolution 3.91 Å). Given that these reconstructions were classified based on a masked region covering only the IASU, the map quality deteriorated with increasing distance from this region, such that the quality was quite poor on the diametrically opposing side of the particle. Reconstructions were subsequently attempted with the larger 3D classes identified in the pH5.6 data prior to the symmetry expansion step. However, resolutions of no better than 5 Å could be obtained for either consensus I1 or symmetry-expanded C1 maps. Furthermore, comparisons of these maps with the model for the “large” class suggested no significant conformational differences beyond rigid body motions of the subunits. Thus, no further models were generated from these 3D classes.

A summary of data collection, processing and analysis is given in Table S2. Structural figures and movies were prepared using ChimeraX ^37^, Blender ^38^ and Molecular Nodes ^39^.

## Data Availability

The structures of the virus-like particles at pH 7.6, pH 6.25, pH 5.9 and pH 5.6, their associated atomic coordinates and raw data, have been deposited into the Electron Microscopy Data Bank (EMDB), the Protein Data Bank (PDB) and the Electron Microscopy Public Image Archive (EMPIAR), with EMDB accession codes EMD-15134, EMD-15209, EMD-15112, EMD-15266, EMD-15348, EMD-15339, EMD-15307, PDB accession codes 8A41, 8A6J, 8A3C, 8ACH, 8AC6 and 8AAY, and EMPIAR accession codes EMPIAR-11065, EMPIAR-11082, EMPIAR-11060 and EMPIAR-11083.

## Acknowledgements

The authors thank Elaine Barclay and Kim Findlay for TEM training and assistance. We thank Matt Byrne and Tom Dendooven for helpful discussions about cryo-EM data processing. We thank Tatiana Domitrovic for discussions and providing the baculovirus plasmid. We thank Todd Yeates, Michael Sawaya and James Sawaya for valuable feedback and comments on the manuscript and for providing helpful support. We thank Albert Bonmassip and Brady Johnston for advice and help with the animations with Blender. At the John Innes Centre, this work was supported by the United Kingdom Biotechnology and Biological Sciences Research Council (BBSRC) Synthetic Biology Research Center “OpenPlant” award (BB/L014130/1), the Institute Strategic Programme Grants “Molecules from Nature—Enhanced Research Capacity” (BBS/E/J/000PR9794) and “Harnessing Biosynthesis for Sustainable Food and Health” (BB/X01097X/1), a capital grant award (BBSRC) to establish Cryo-EM capability at the John Innes Centre and the John Innes Foundation. The cryo-EM microscopes at the Astbury Biostructure Laboratory (ABSL) were funded by the University of Leeds (UoL ABSL award) and Wellcome Trust (108466/Z/15/Z). Use of the Stanford Synchrotron Radiation Lightsource, SLAC National Accelerator Laboratory, is supported by the U.S. Department of Energy, Office of Science, Office of Basic Energy Sciences under Contract No. DE-AC02-76SF00515. The SSRL Structural Molecular Biology Program is supported by the DOE Office of Biological and Environmental Research, and by the National Institutes of Health, National Institute of General Medical Sciences (P30GM133894). The contents of this publication are solely the responsibility of the authors and do not necessarily represent the official views of NIGMS or NIH. The Pilatus detector at beamline 4-2 at SSRL was funded under National Institutes of Health Grant S10OD021512.

