## Supplementary Material for "Unravelling the Maturation Pathway of a Eukaryotic Virus through Cryo-EM"

Castells-Graells, *et al.*

### Supplementary Tables:

**Table S1. SAXS data collection and analysis**

| | |
|---|---|
| Data collection | |
| Beamline | SSRL BL4-2 |
| Type of experiment | Equilibrium SAXS |
| Beam defining slits size | 0.3 x 0.3 mm |
| Sample-Detector distance | 1.7 m |
| Beam energy | 11.0 keV |
| Beam current | 500 mA |
| Exposure time per frame | 3 sec |
| Detector | Pilatus 3 X 1M |
| Frames per data set | 10 |
| Sample cell size | 1.5 mm in diameter |
| Temperature | 293 K |
| Sample volume | 30 μL |
| Sample concentration | 0.94 mg/ml |
| *q* range* | 0.0066 - 0.51 |
| Software | |
| Data reduction | SASTool/SAXSPipe |
| *P*(r) estimation | Primus/GNOM |

* $q = 4\pi\sin(\theta)/\lambda$, where $2\theta$ is the scattering angle.

### Table S2. Summary of cryo-EM data collection, processing and analysis

| Data set | Procapsid (pH 7.6) | pH6.25 | pH5.9 | pH5.6 consensus | pH5.6 large | pH5.6 medium | pH5.6 small |
|---|---|---|---|---|---|---|---|
| **Data collection and processing** | | | | | | | |
| Microscope | Titan Krios | Titan Krios | Titan Krios | Titan Krios | - | - | - |
| Detector (mode) | Falcon III (integrating) | Falcon III (integrating) | Falcon III (integrating) | Falcon III (integrating) | - | - | - |
| Magnification | 75,000 x | 75,000 x | 75,000 x | 75,000 x | - | - | - |
| Magnified pixel size (Å) | 1.065 | 1.065 | 1.065 | 1.065 | - | - | - |
| Voltage (kV) | 300 | 300 | 300 | 300 | - | - | - |
| Total dose ($e^-$/Å$^2$) | 88.5 | 50.4 | 46.0 | 48.7/50.4 | - | - | - |
| Defocus range (μm) | -0.7 to -2.5 | -0.8 to -2.2 | -0.7 to -2.2 | -0.8 to -2.5 | - | - | - |
| Movies collected | 6,926 | 10,528 | 10,010 | 16,478 | - | - | - |
| Particle images | 33,881 | 114,943 | 38,792 | 15,977 | 96,960[†] | 264,917[†] | 353,898[†] |
| Symmetry imposed | Icosahedral (I1) | Icosahedral (I1) | Icosahedral (I1) | Icosahedral (I1) | None (C1) | None (C1) | None (C1) |
| FSC threshold | 0.143 | 0.143 | 0.143 | 0.143 | 0.143 | 0.143 | 0.143 |
| Map resolution (Å) | 4.88 | 4.80 | 3.92 | 3.39 | 3.91 | 3.63 | 3.39 |
| Map resolution range (Å) | 4.22 – 9.78 | 4.26 – 9.24 | 3.49 – 7.73 | 3.18 – 6.93 | 3.48 – 10.17 | 3.39 – 7.69 | 3.10 – 6.49 |
| **Refinement*** | | | | | | | |
| Initial model used (PDB code) | 8A3C | 8A3C | 7ANM | - | 8AC6 | 8AAY | 7ANM |
| Map sharpening *B* factor (Å$^2$) | -260.5 | -432.6 | -233.0 | -149.6 | -50.0 | -100 | -100 |
| Model composition (ASU) | Chains A-D | Chains A-D | Chains A-D | - | Chains A-D | Chains A-D | Chains A-D |
| Non-hydrogen atoms | 17,464 | 17,400 | 17,197 | - | 16,450 | 16,373 | 16,702 |
| Protein residues | 2,288 | 2,272 | 2,246 | - | 2,150 | 2,142 | 2,180 |
| R.m.s. deviations | | | | | | | |
| Bond lengths (Å) | 0.004 | 0.004 | 0.004 | - | 0.002 | 0.003 | 0.006 |
| Bond angles (°) | 0.93 | 0.93 | 0.93 | - | 0.47 | 0.46 | 1.00 |
| Model diameter (full particle; Å) | 470 | 460 | 455 | - | 430 | 425 | 415 |
| **Validation** | | | | | | | |
| Molprobity score | 2.53 | 2.18 | 2.44 | - | 2.43 | 2.23 | 1.91 |
| Clashscore | 12.7 | 8.8 | 9.54 | - | 9.26 | 7.31 | 6.16 |
| Poor rotamers (%) | 5.24 | 3.00 | 5.79 | - | 6.01 | 4.69 | 2.88 |

| | | | | | | | |
|---|---|---|---|---|---|---|---|
| Ramachandran plot: | | | | | | | |
| Favoured (%) | 94.6 | 95.1 | 94.9 | - | 95.0 | 95.3 | 96.5 |
| Allowed (%) | 5.4 | 4.9 | 5.1 | - | 5.0 | 4.6 | 3.4 |
| Disallowed (%) | 0.0 | 0.0 | 0.0 | - | 0.0 | 0.1 | 0.1 |
| Ramachandran Z-score | -0.66 ± 0.19 | -0.08 ± 0.19 | -1.16 ± 0.19 | - | -0.02 ± 0.19 | -0.50 ± 0.19 | -0.61 ± 0.18 |
| Fit to map ($CC_{mask}$) | 0.83 | 0.75 | 0.77 | - | 0.75 | 0.77 | 0.87 |
| **Accession codes** | | | | | | | |
| EMPIAR (dataset) | 11065 | 11082 | 11060 | 11083 | 11083 | 11083 | 11083 |
| EMDB (maps) | 15134 | 15209 | 15112 | 15266 | 15348 | 15339 | 15307 |
| PDB (model) | 8A41 | 8A6J | 8A3C | - | 8ACH | 8AC6 | 8AAY |

*Refinement statistics refer to IASU, not the whole particle.

†Figures are for the symmetry expanded set and therefore refer to the number of IASUs used in the subsequent processing steps.

**Supplementary Figures:**

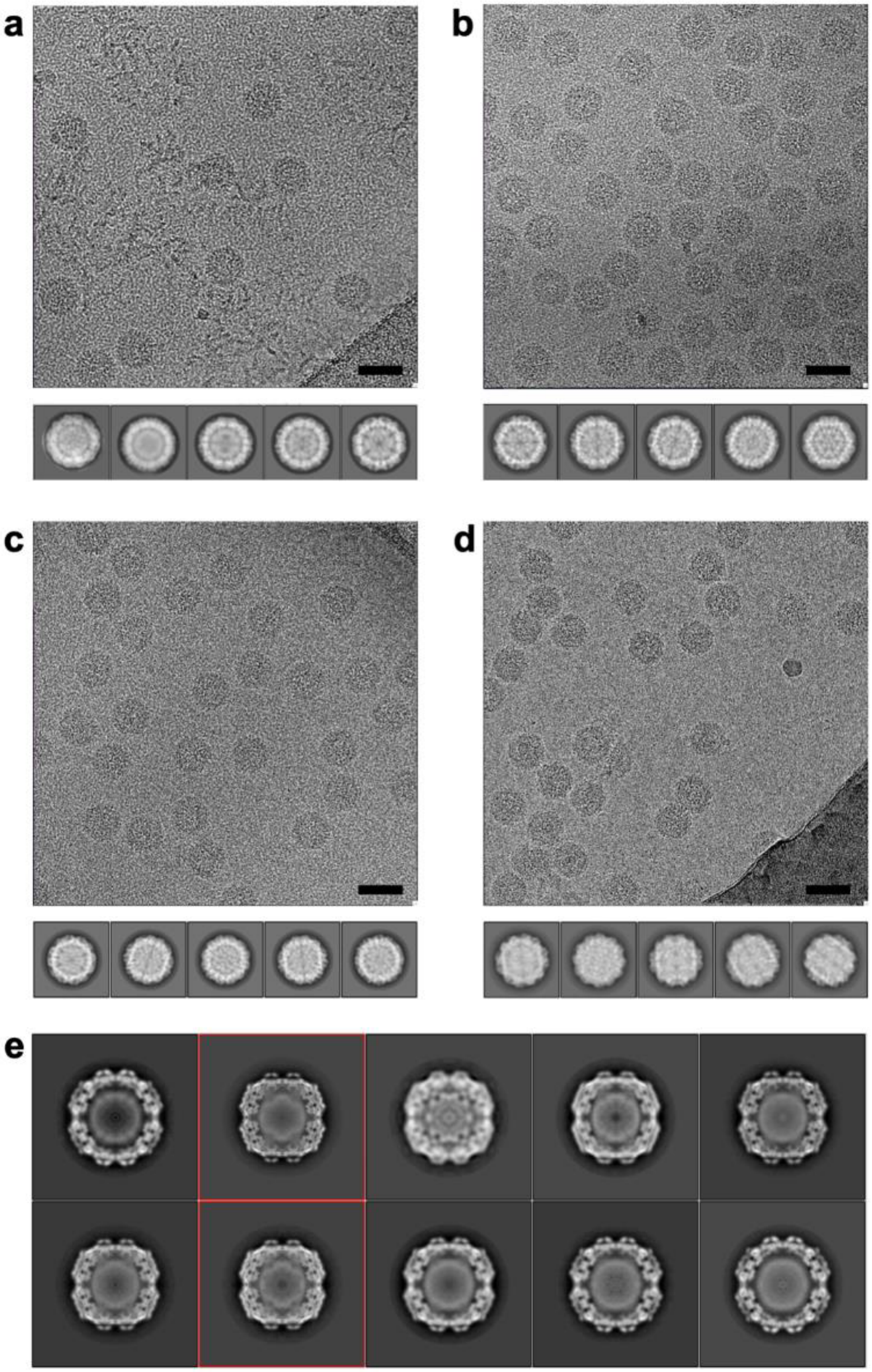

**Figure S1.** Cryo-EM data collection and preliminary processing. Motion-corrected movies (MOTIONCOR2) showing **(a)** procapsid at pH 7.6, **(b)** pH 6.25 intermediate, **(c)** pH 5.9 intermediate and **(d)** pH 5.6 intermediate NωV particles imaged in vitreous ice. The scale bar = 50 nm. Below each image are 2D class averages (Relion) obtained from these data sets (not to same scale). Panel **(e)** shows 3D classes for the pH 5.6 intermediate processed with icosahedral symmetry (I1). The two smallest classes highlighted in red were combined and taken forward for symmetry expansion and further 3D classification.

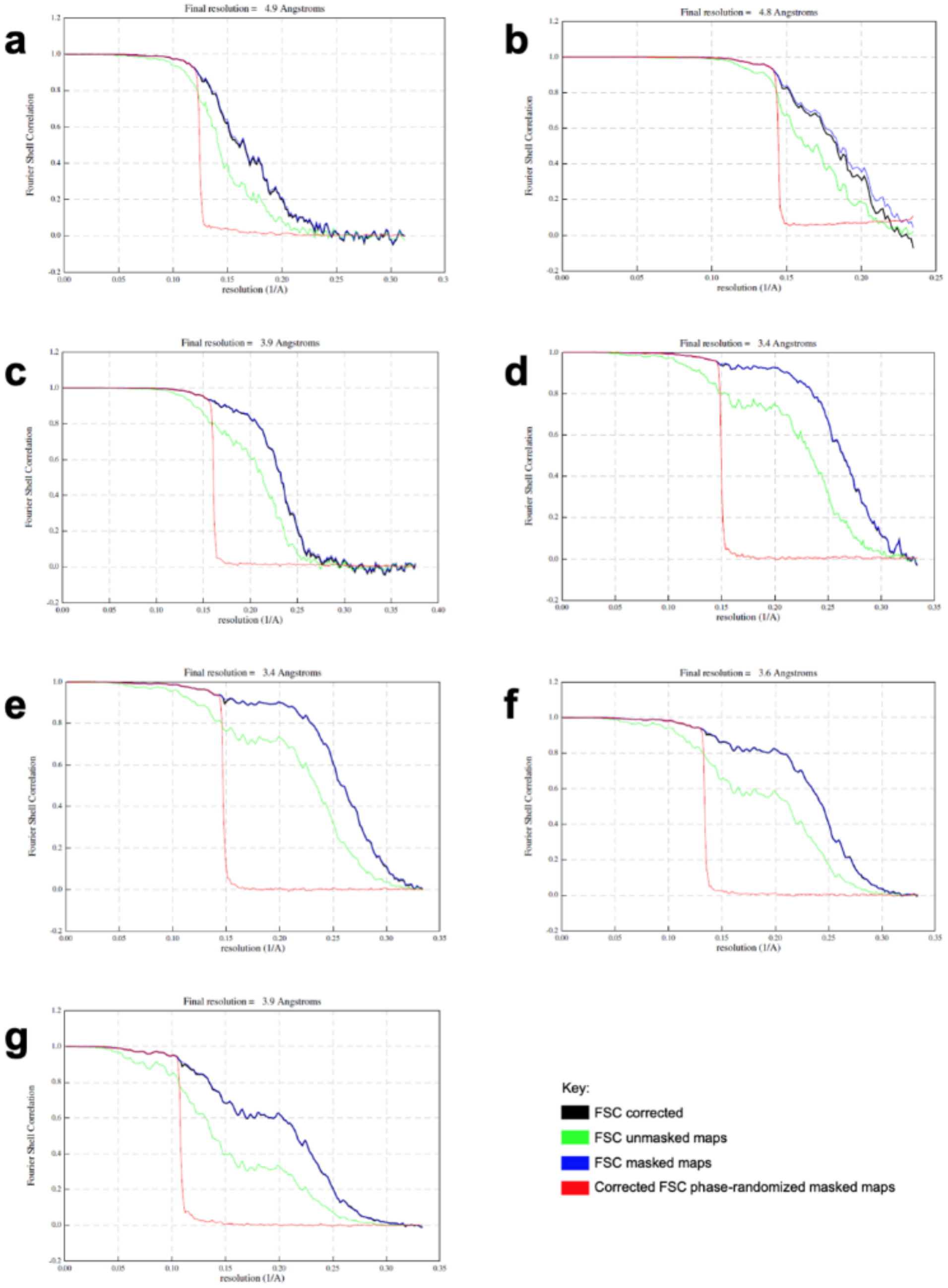

**Figure S2.** Cryo-EM postprocessing. Fourier Shell Correlation curves generated in the final postprocessing step of Relion for **(a)** procapsid, **(b)** pH 6.25 intermediate and **(c)** pH 5.9 intermediate. **(d)** Shows the curve for pH 5.6 consensus map and **(e-f)** show the curves for the small, medium and large symmetry-expanded reconstructions. The resolutions were assessed using FSC = 0.143.

**Supplementary Movies:**

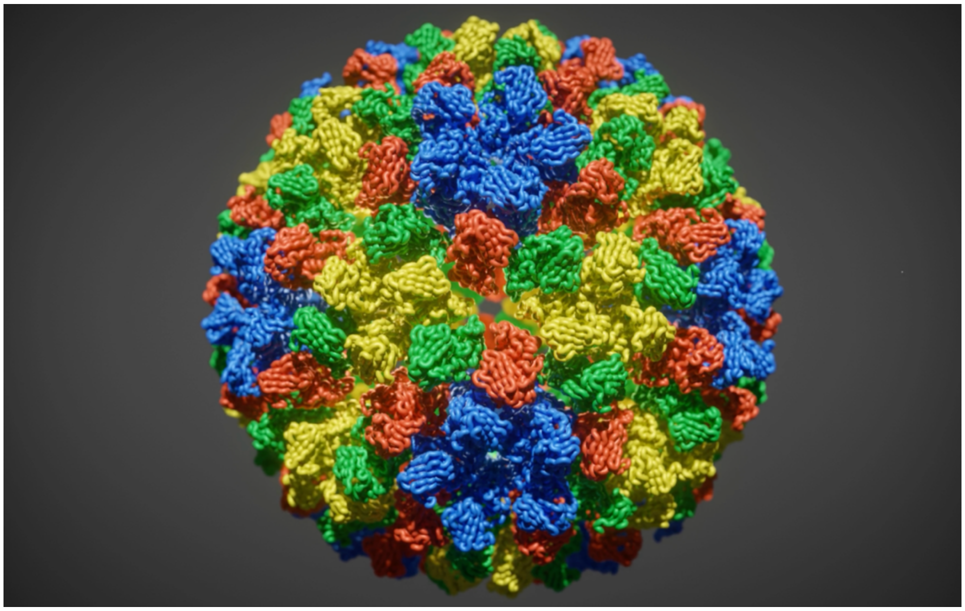

**Movie S1.** NωV maturation process. Ribbon representation of the NωV particle morphing between the different intermediates in the maturation process, from procapsid to capsid. Subunits are color coded A (blue), B (red), C (green) and D (yellow).

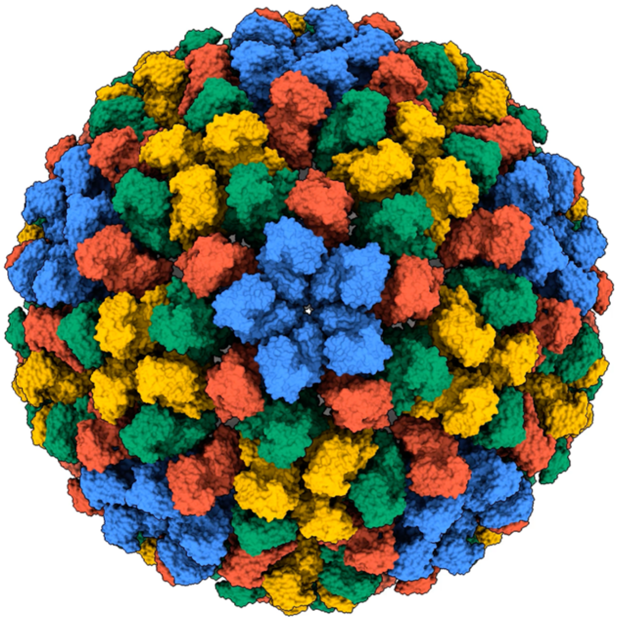

**Movie S2.** Surface view down a 5-fold axis of the NωV particle morphing between the different intermediates in the maturation process.

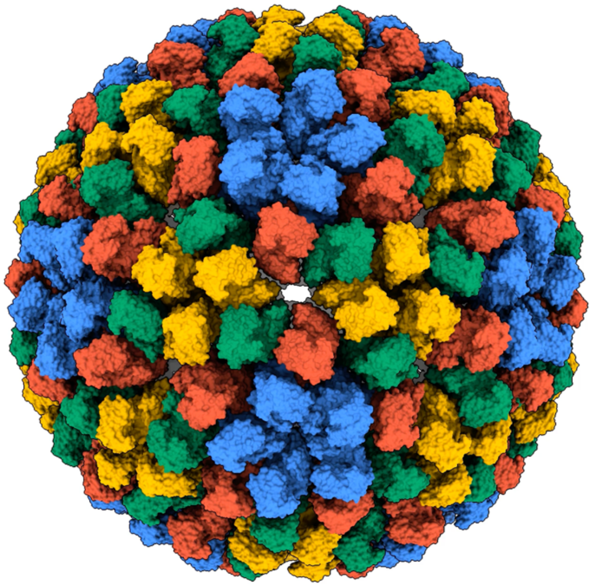

**Movie S3.** Surface view down a quasi-6-fold (icosahedral 2-fold) axis of the NωV particle morphing between the different intermediates in the maturation process.

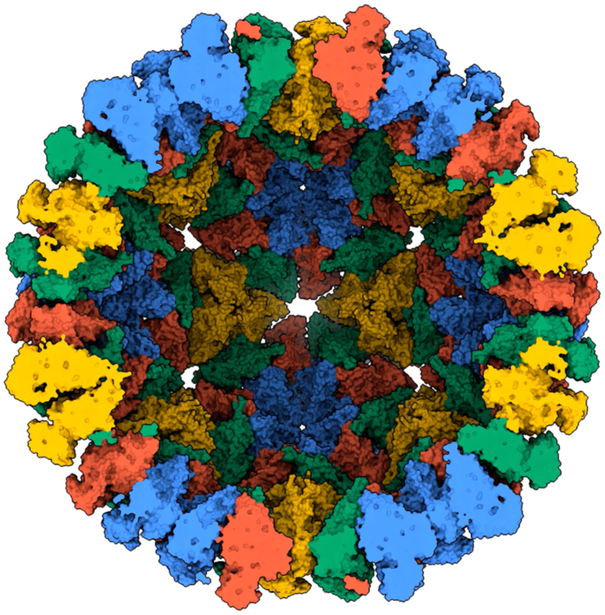

**Movie S4.** Surface representation down a quasi-6-fold axis of a half section of a NωV particle morphing between the different intermediates in the maturation process.

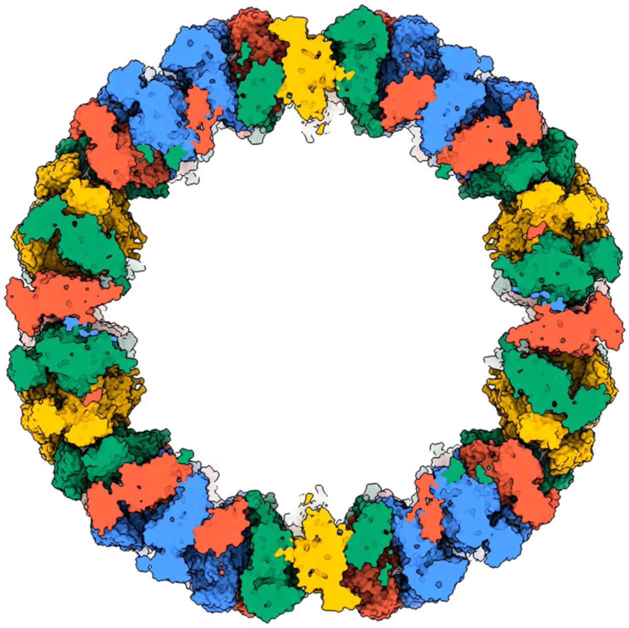

**Movie S5:** Surface representation down a quasi-6-fold axis of a middle section of a NωV particle morphing between the different intermediates in the maturation process.

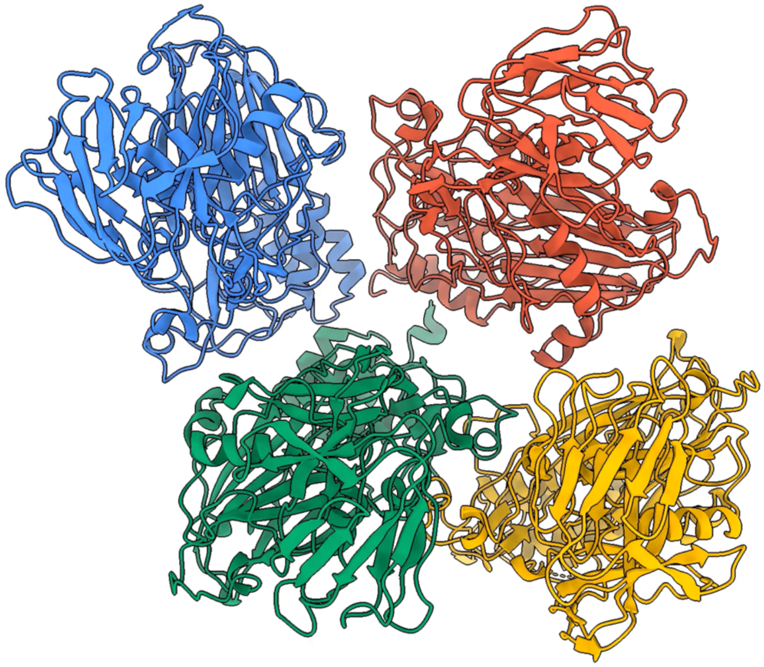

**Movie S6:** Ribbon representation of the NωV ASU morphing between the different intermediates in the maturation process.

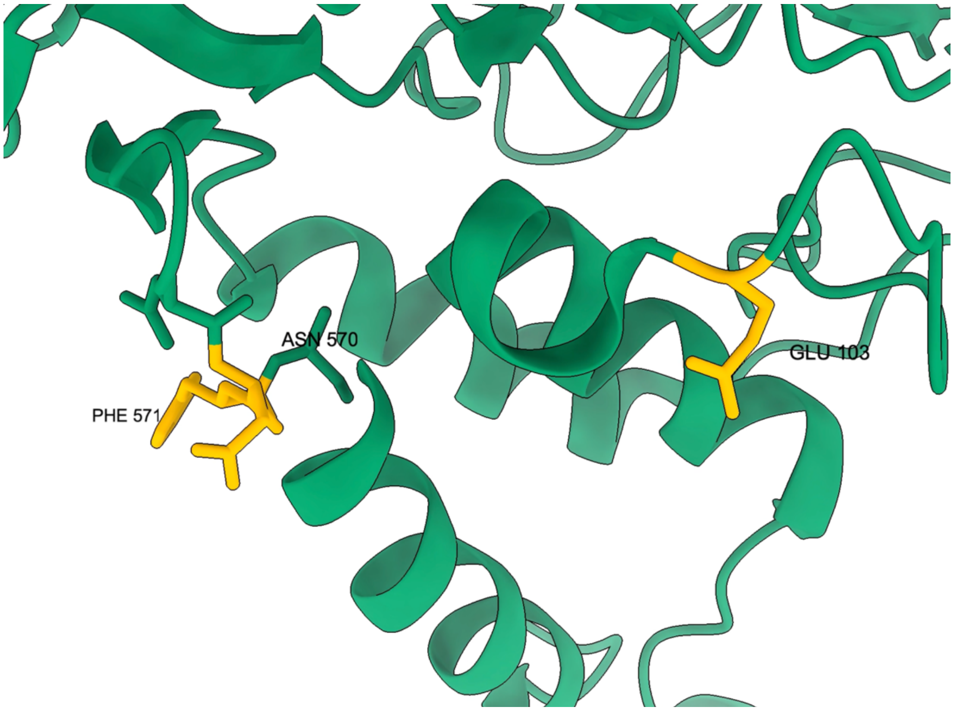

**Movie S7:** Ribbon representation of the cleavage site of the NωV subunit C morphing between different intermediates in the maturation process, from procapsid to capsid.
